# Head-related transfer functions of rabbits within the front horizontal plane

**DOI:** 10.1101/2023.09.15.557943

**Authors:** Mitchell L. Day

## Abstract

The head-related transfer function (HRTF) is the direction-dependent acoustic filtering by the head that occurs between a source signal in free-field space and the signal at the tympanic membrane. HRTFs contain information on sound source location via interaural differences of their magnitude or phase spectra and via the shapes of their magnitude spectra. The present study characterized HRTFs for source locations in the front horizontal plane for nine rabbits, which are a species commonly used in studies of the central auditory system. HRTF magnitude spectra shared several features across individuals, including a broad spectral peak at 2.6 kHz that increased gain by 12 to 23 dB depending on source azimuth; and a notch at 7.6 kHz and peak at 9.8 kHz visible for most azimuths. Overall, frequencies above 4 kHz were amplified for sources ipsilateral to the ear and progressively attenuated for frontal and contralateral azimuths. The slope of the magnitude spectrum between 3 and 5 kHz was found to be an unambiguous monaural cue for source azimuths ipsilateral to the ear. Average interaural level difference (ILD) between 5 and 16 kHz varied monotonically with azimuth over ±31 dB despite a relatively small head size. Interaural time differences (ITDs) at 0.5 kHz and 1.5 kHz also varied monotonically with azimuth over ±358 μs and ±260 μs, respectively. Remeasurement of HRTFs after pinna removal revealed that the large pinnae of rabbits were responsible for all spectral peaks and notches in magnitude spectra and were the main contribution to high-frequency ILDs, whereas the rest of the head was the main contribution to ITDs and low-frequency ILDs. Lastly, inter-individual differences in magnitude spectra were found to be small enough that deviations of individual HRTFs from an average HRTF were comparable in size to measurement error. Therefore, the average HRTF may be acceptable for use in neural or behavioral studies of rabbits implementing virtual acoustic space when measurement of individualized HRTFs is not possible.

## 1. Introduction

The sound pressure wave reaching the tympanic membrane is not the same as that of its source in free-field space. Along the path from source to tympanic membrane, sound pressure is linearly transformed by reflections off of and diffraction around the head and body; constructive and destructive interference due to reflections off of features of the pinna; and resonance in open cavities of the outer ear. This transformation of pressure magnitude and phase, termed the head-related transfer function (HRTF), is frequency- and direction-dependent; therefore, sound pressure will be amplified at some frequencies and attenuated at others depending on the direction of the source. The set of HRTFs across locations in space contain the cues that are known to be perceptually relevant to the determination of the direction of a sound source: interaural time and level differences (ITDs and ILDs) for determination of source azimuth and shape of the magnitude spectrum for determination of source elevation (Middlebrooks and Green, 1991).

Knowledge of the HRTFs of a species is scientifically useful for at least two reasons. First, HRTFs contain information on how sound waveforms that reach the tympanic membrane are differentially weighted across frequency for that species. For example, many mammals exhibit a broad and largely non-directional peak at a particular frequency in the HRTF magnitude spectrum, attributed to acoustic resonance within the ear canal (Shaw and Teranishi, 1968). Therefore, frequency components of the source signal at and around the resonant frequency for that species are emphasized in the signal input to the organism. Second, HRTFs contain information on the range of binaural cues and the characteristics of monaural spectral cues available to a species for sound localization. For example, the range of ITDs and ILDs across spatial locations varies across species and across frequency within a given species. This information is important when interpreting neural or behavioral responses of a given species to sounds presented over headphones where it is possible to create ITDs or ILDs outside of the range for free-field sources.

HRTFs have been measured in humans, many other mammalian species, and some non-mammalian species. In the present study, we report HRTFs of rabbits—a species known for their relatively long pinnae. Rabbits have a hearing range that largely overlaps with that of humans (Heffner and Masterton, 1980) and are commonly used in neurophysiological and behavioral studies of the auditory system (Barzelay et al., 2023; Carney et al., 2014; Fan et al., 2022; Haragopal et al., 2020; Kim et al., 2020; Kuwada et al., 2015; Wagner et al., 2022; Zhai et al., 2020). There has been one previous study of rabbit HRTFs (Kim et al., 2010). Like the present study, Kim et al. (2010) measured HRTFs of the Dutch-belted strain of rabbits in the front horizontal plane and reported magnitude spectra, ILD spectra, and ITD spectra. However, unlike the present study, their study focused on changes of the HRTF with distance of the sound source and comparisons to a spherical head model. Our data complement that of Kim et al. (2010) by characterizing features of magnitude spectra shared across rabbits and examining the dependence of those features on source azimuth. While Kim et al. (2010) reported data from two rabbits and made no comparisons between them, we examined inter-individual differences across nine rabbits. Additionally, we assessed the potential utility of monaural spectral cues for determination of source azimuth and investigated the contribution of the pinnae to binaural cues and features of magnitude spectra.

HRTFs can be used to filter sound waveforms presented over earphones so that they are perceived as indistinguishable from an externalized, free-field source (Kulkarni and Colburn, 1998). This “virtual acoustic space” technique is advantageous because it allows presentation of free-field-like sounds monaurally or with augmented localization cues in order to dissect function in neurophysiological or psychophysical experiments (Dorkoski et al., 2020). Experiments utilizing virtual acoustic space are difficult because a subject’s or animal’s own HRTFs must be used to minimize experimental error, assuming HRTFs are significantly different across individuals. To assess this assumption for rabbits, we quantified inter-individual differences of HRTFs and differences between individual HRTFs and an average HRTF. Our results may be of use to researchers who wish to perform neurophysiological or behavioral experiments on rabbits using virtual acoustic space, but who do not have the capability of measuring individualized HRTFs.

## 2. Methods

### 2.1. Animal preparation

A total of nine Dutch-belted rabbits were used in the study, including four females and five males. All came from the same breeding facility (Envigo). Six rabbits shared the same birthdate and were likely related, while the other three shared a birthdate a year later. Rabbits weighed between 1.91 and 2.28 kg and were of ages between 35 and 88 weeks at the time of measurement. Three rabbits had an aluminum headbar and electrode microdrive mounted to the top of the skull for use in neurophysiological experiments in a separate study. The microdrive and drive holder portion of the headbar (together, 2.7 cm × 2.2 cm × 2.8 cm) were positioned directly above the skull, and the headbar (5.2 cm × 1 cm × 1 cm) projected anteriorly. All animal procedures were approved by the Ohio University Institutional Animal Care and Use Committee.

All rabbits were used for additional studies. Therefore, acoustic measurements usually preceded another anesthetized procedure, such as measurement of auditory brainstem responses or implantation of the headbar. Acoustic measurements were completed in less than 10 minutes. Rabbits were anesthetized with either xylazine (6 mg/kg s.c.) and ketamine (35 mg/kg i.m.), or xylazine (6 mg/kg s.c.), acepromazine (1 mg/kg s.c.) and ketamine (44 mg/kg i.m.). Glycopyrrolate (0.01 mg/kg s.c.) or atropine (0.25 mg/kg s.c.) was administered to reduce mucosal secretions and prevent tracheal blockage. Measurements proceeded when rabbits lacked a foot-pinch reflex. The anesthetized rabbit was laid down in prone position inside a sound-attenuating chamber and monitored via webcam during measurement. Following measurement, or the subsequent procedure, rabbits recovered from anesthesia on a warm heating pad while receiving pure oxygen via mask, and heart rate and blood oxygenation level were monitored. One rabbit was euthanized (pentobarbital, 100 mg/kg i.v.) and refrigerated in the same prone position prior to acoustic measurement in order to study the effect of pinna removal. After pinna removal, only about 1 mm of cartilaginous ring was left protruding from the bony ear canal on each side.

### 2.2. Acoustic measurement

Acoustic measurements took place inside a double-walled, sound-attenuating chamber (Acoustic Systems) of interior dimensions 2.13 m wide × 2.84 m deep × 2.31 m high. Walls and ceiling were lined with 5-cm, sound-absorbing foam (SONEX with acoustic coating, Pinta Acoustic). Inside the chamber was a custom-made, metal, 1-m-radius, semicircular array of 25 full-range speakers (Rockford Fosgate R14X2) spaced 7.5° apart. The ends of the array were mounted on swivels so that the array could be rotated to different elevations. The array could be fixed to specific elevations in the top hemisphere from 0° (front) to 180° (back) in 7.5° increments by pinning it with heavy pins into pre-drilled holes in fixed, metal plates at the array ends. The whole array could be translated vertically up to 1.83 m above the floor by sliding along four metal posts (two on each side). However, in the present study, vertical height of the horizontal plane was kept at approximately the height of the rabbit’s ears. Before measurements, metal posts, metal plates, and the area of floor inside the circle of the array were covered in sound-absorbing foam.

Stimulus presentation and data acquisition were controlled by custom code written with LabVIEW software (National Instruments) running on a PC. Stimuli were created digitally, converted to analog signals at 48,828.125-Hz sampling rate with 24-bit resolution (Tucker-Davis Technologies [TDT] RZ6), amplified (TDT RZ6), and switched to drive each of the 25 speakers in the array via two daisy-chained multiplexers (TDT PM2R). The stimulus was a frequency-sweeping chirp of 20-ms duration presented at a fixed attenuation level and repeated continuously 512 times. Its magnitude spectrum was flat and phase was chosen to minimize peak factor (Schroeder, 1970). The attenuation level was chosen to reduce the amplitude of echoes in the measured acoustic signal while keeping the signal above the noise floor at all frequencies. Acoustic signals were measured with a probe-tube microphone (Etymotic ER-7C) and digitized at 48,828.125-Hz sampling rate with 24-bit resolution (TDT RZ6).

Speaker-only acoustic measurements in the absence of the rabbit were made for each speaker location by dangling the probe-tube microphone so that the end of the probe tube was in the center of the speaker array at the approximate height of a rabbit’s ears. In-ear acoustic measurements were made with the end of the probe tube approximately inside the entrance of the ear canal. The flexible probe-tube end was glued with cyanoacrylate midway inside a cylindrical shell of heat-shrink tubing of 5 mm length, 5 mm diameter, and less than 1 mm thickness. The shell with probe tube was inserted into the concha of the pinna with tweezers until a snug fit was attained. For in-ear measurements, each rabbit was laid down prone on the sound-absorbing foam with the midway-between-ears position of the head at the center of the speaker array, snout pointed toward the central speaker (0° azimuth), and pinna in a natural, relaxed backward position (Fig. 1). Speaker-only and in-ear measurements were made for speaker locations in the front horizontal plane at azimuths from 90° left (L) to 90° right (R) in 15° steps.

**Fig. 1.**
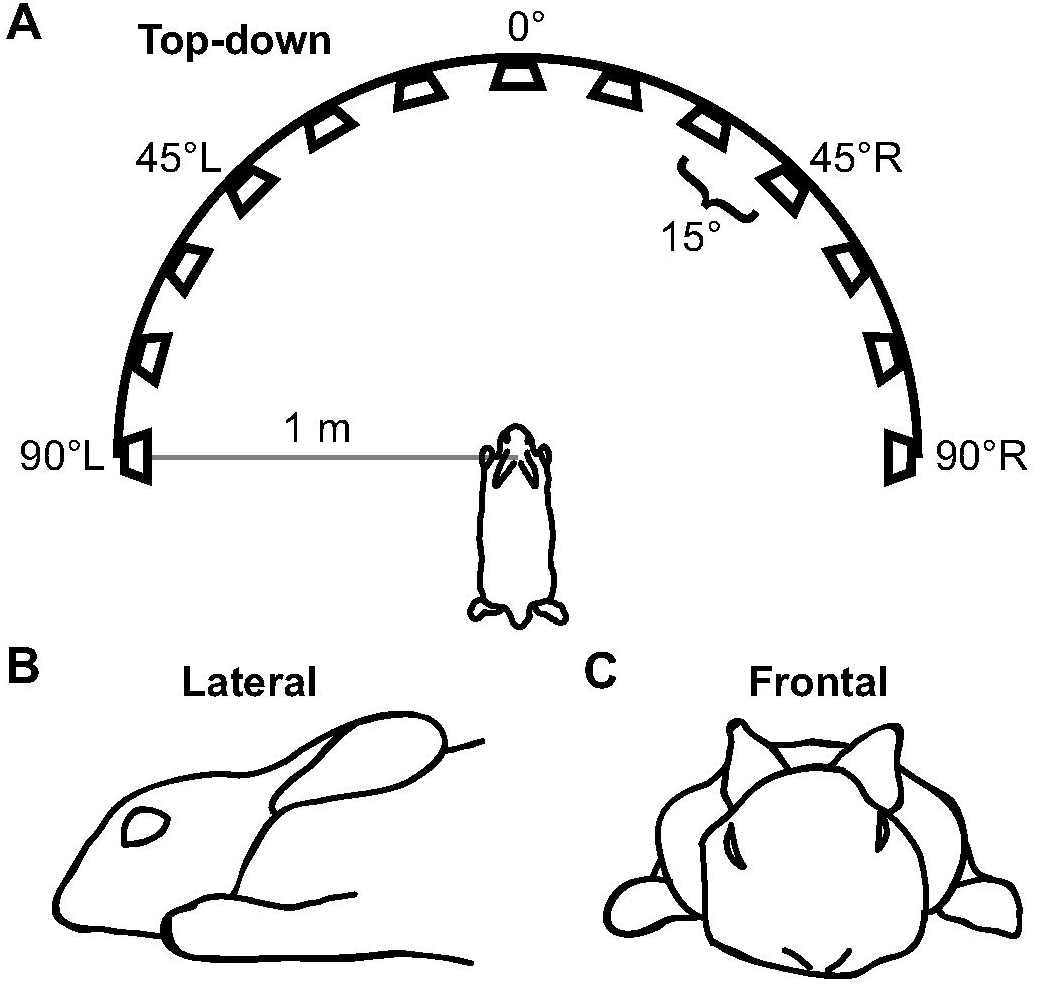
Top-down (A), lateral (B), and frontal (C) views of the rabbit within the speaker array during HRTF measurement. Drawings are tracings of photographs.

### 2.3. Signal processing

All signal processing, data analysis, and statistical tests were performed using MATLAB software (Mathworks). Computation of head-related transfer functions (HRTFs) followed standard linear filtering principles (Mehrgardt and Mellert, 1977). For a given speaker location, the speaker-only impulse response is the circular cross-correlation of the stimulus waveform and the average speaker-only response waveform. This computation was made in the frequency domain as *x*[*n*] = *F*^-1^{*S*\*(*w*)*R_Speaker_*(*w*)}, where *x*[*n*] is the speaker-only impulse response, *n* is the time sample, *F*^-1^ represents the inverse Fourier transform, *S\**(ω) is the complex conjugate of the Fourier transform of the stimulus waveform, *R_speaker_*(ω) is the Fourier transform of the average speaker-only response waveform, and ω is frequency in radians/sec. Similarly, the in-ear impulse response, *y*[*n*], for a given speaker location and ear was *y*[*n*] = *F*^-1^{*S*\*(*w*)*R_ear_*(*w*)}, where *R_ear_*(ω) is the Fourier transform of the average in-ear response waveform. Impulse responses were resampled at 50-kHz sampling rate. In-ear impulse responses could display prominent echoes, particularly for speaker locations contralateral to the ear (Fig. 2A). The effect of echoes was mitigated by windowing impulse responses with a Hann window of 5-ms duration and centered at 5.5 ms, which corresponded to the time of the peak of the impulse response for the speaker located at 0° azimuth.

**Fig. 2.**
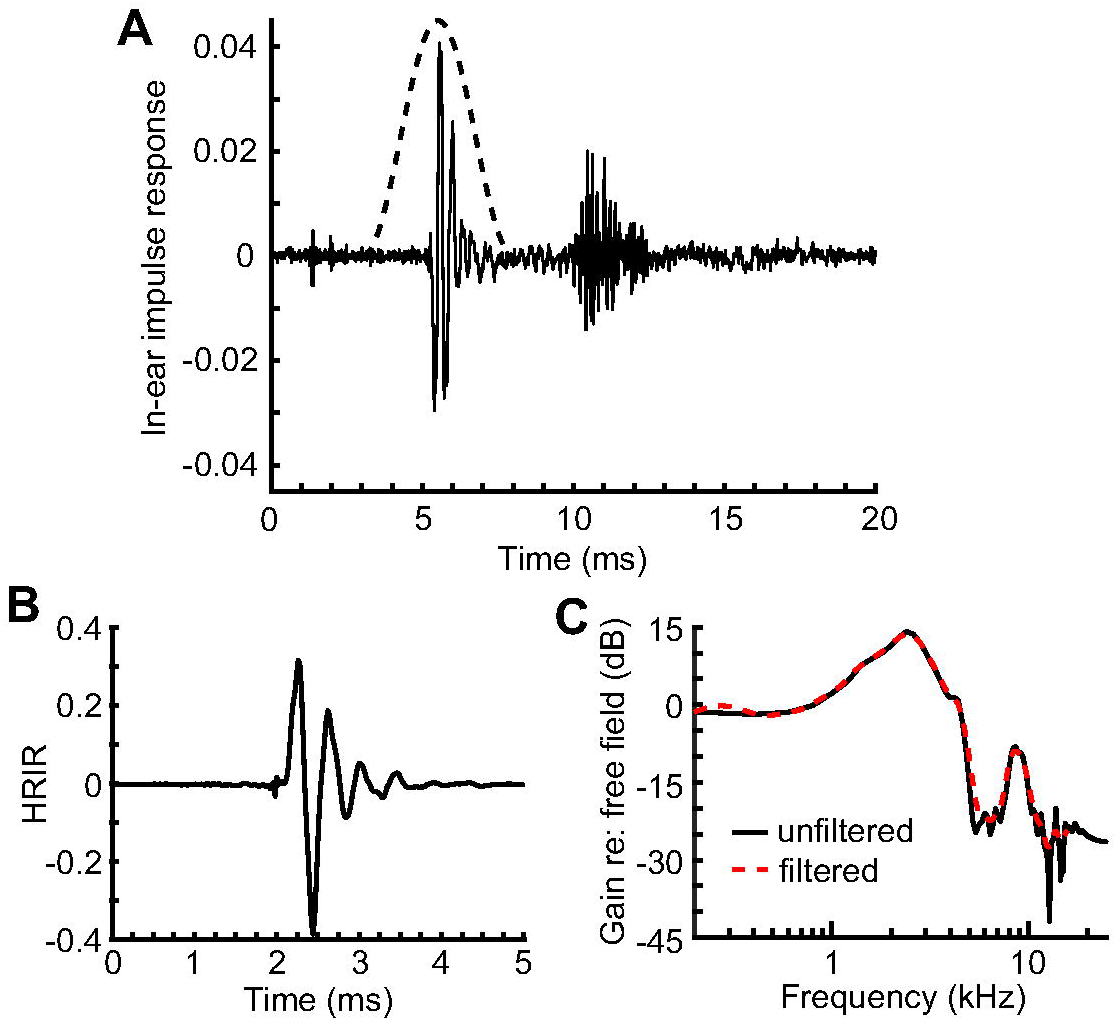
Signal processing. (A) In-ear impulse response showing an initial component from the direct sound wave isolated by a Hann window (dashed), and a delayed component from echoes. (B) HRIR after low-pass filtering. (C) Magnitude spectrum of the HRTF before (black solid) and after (red dashed) passing through a filter bank. All panels show data from the same ear for a source at 45°C.

The in-ear impulse response includes not only filtering properties of the head but also filtering due to the transfer functions of the speaker and microphone. To isolate the head-related component, the in-ear impulse response is divided by the speaker-only impulse response in the frequency domain. Specifically, the head-related impulse response (HRIR) was HRIR(*w*) = *F*^-1^{*Y*(*w*)/*X*(*w*)}, where *Y*(ω) and *X*(ω) are the Fourier transforms of the in-ear and speaker-only impulse responses, respectively. Before performing the inverse Fourier transform operation in the preceding equation, the DC component of the argument was set to zero to prevent a large DC shift in the HRIR that could occur due to dividing by a very small number.

The frequency response of the probe-tube microphone was flat up to 10 kHz, smoothly decreased between 10 and 16 kHz, but was unreliable above 16 kHz. Therefore, as a final step, we low-pass filtered the HRIR at a cutoff frequency of 16 kHz using a fourth-order Butterworth filter (Fig. 2B).

### 2.4. Data analysis

The HRTF is the Fourier transform of the HRIR. Magnitude spectra of HRTFs often displayed very narrow peaks and notches at high frequencies (Fig. 2C). Such fluctuations are of likely little physiological relevance due to the relatively large bandwidth of cochlear filtering as compared to the widths of the narrow peaks and notches. In order to characterize HRTF magnitude spectra based on physiologically relevant features, each HRIR was first passed through a bank of fourth-order gammatone filters to approximate filtering by the basilar membrane of the cochlea (Zhang et al., 2001). The impulse response of each gammatone filter was *g*(*t*) = (1/*β*)*t*^γ-1^ exp (-*t*/τ)cos (2*πf_CF_t*), where *t* is time from 0 to 40 ms, γ is the filter order, τ is the filter time constant, and *f_CF_* is the characteristic frequency of the filter. The normalization constant, β, was chosen so that the total energy of the filter was one. The filter time constant was defined as τ = (2*Q*_10_)/(2*πf_CF_*), where *Q*_10_ is the ratio of the *f_CF_* of an auditory nerve fiber to the bandwidth of the fiber’s response measured 10 dB above threshold (Zhang et al., 2001). *Q*_10_ values were taken from a piecewise-linear fit of data from rabbit auditory nerve fibers (Borg et al., 1988): *Q*_10_(*f_CF_*) =2 for *f_CF_* ≤ 1.5 kHz; *Q*_10_(*f_CF_*) = 4.814 log_10_ *f_CF_* + 1.152 for 1.5 kHz < *f_CF_* ≤ 8 kHz; and *Q*_10_(*f_CF_*) = 5.5 for *f_CF_* > 8 kHz, with *f_CF_* in units of kHz. A bank of filters was created with *f_CF_*values from 0.2 to 16 kHz in 0.05-oct steps, where the logarithmic spacing of characteristic frequencies approximated that along the basilar membrane. Filtering of the HRIR was performed in the frequency domain: HRTF_filtered_(ω) = G(ω)HRTF(ω), where *G*(ω) is the Fourier transform of the gammatone impulse response. The output of the filter bank was characterized by the square root of the energy of each filtered HRTF: [(1⁄*N*) ∑_ω_|HRTF_filtered_(ω)|^2^]^1/2^, where *N* is the total number of frequency samples. The filter bank had the desired effect of smoothing out fluctuations in HRTF magnitude spectra (Fig. 2C). All analyses of magnitude spectra and ILD were performed on the output of the filter bank; for simplicity, the output and *f_CF_* are referred to below as “gain” and “frequency”, respectively.

Prominent mid-frequency peaks were apparent in magnitude spectra at all azimuths, but prominent high-frequency notches and peaks were only apparent at some azimuths. A high-frequency notch was defined as a minimum in the magnitude spectrum between 4 and 9.1 kHz with gain rising continuously by at least 5 dB for frequencies above and below the minimum before reversing. Similarly, a high-frequency peak was defined as a maximum in the magnitude spectrum between 8.7 and 14 kHz with gain falling continuously by at least 5 dB for frequencies above and below the maximum before reversing.

The utility of monaural spectral cues for the use of localization was examined by comparing average gain within seven neighboring frequency bands. Each band had a width of 0.38 oct, which was equal to the frequency distance between the high-frequency notch and peak. Frequency borders of the bands were: (*i*) 1.8–2.3 kHz, (*ii*) 2.3–3.0 kHz, (*iii*) 3.0–3.9 kHz, (*iv*) 3.9–5.1 kHz, (*v*) 5.1–6.6 kHz, (*vi*) 6.6–8.6 kHz, and (*vii*) 8.6–11.2 kHz. Bands *vi* and *vii* were centered on the high-frequency notch and peak, respectively. Average gain was computed as the average of decibel gain values at frequency components within the band.

The ILD spectrum was computed as either right-ear log-magnitude spectrum minus left-ear log-magnitude spectrum for the same location or ipsilateral-ear log-magnitude spectrum minus contralateral-ear log-magnitude spectrum for the same location depending on whether azimuth was expressed as left/right or ipsilateral/contralateral. Average ILD within a frequency band was computed as the average of decibel ILD values at frequency components within the band. The ITD spectrum was computed as right-ear phase spectrum minus left-ear phase spectrum for the same location, divided by frequency. Phase spectra were computed from the original, unfiltered HRTFs and unwrapped so that phase decreased with frequency.

Lastly, a composite average HRTF across spatial locations was constructed using different averaging methods for the magnitude and phase components: HRTF_avg_(ω) = *r*(ω)*e^iφ^*^(ω)^ where *r*(ω) and φ(ω) are the magnitude and phase, respectively. The magnitude of the average HRTF was computed by averaging log magnitudes:

*r*(ω) = exp((1/*M*) ∑_θ_ log|HRTF_θ_(ω)|) where θ is spatial location and *M* is the total number of locations. The phase of the average HRTF was computed by directly averaging HRTFs: φ(ω) = ((1/*M*)) ∑_θ_ HRTF_θ_(ω)) where < is the angle function.

### 2.5. Statistical tests

Statistical analyses were performed to test for an effect of sex, mass, or age on values from magnitude, ILD and ITD spectra. Sex differences were assessed with a *t*-test if values were sufficiently normally distributed (p ≥ 0.05, Jarque-Bera test), or a Mann-Whitney *U* test if not. Effect size of sex differences was reported as the d-prime (i.e., absolute difference of means divided by the average standard deviation). Correlations between values and mass or age were assessed with Pearson’s linear correlation coefficient if both variables were sufficiently normally distributed, or Spearman’s rho if not.

## 3. Results

### 3.1. General features of HRTF magnitude spectra

We measured HRTFs for spatial locations within the front horizontal plane in the left and right ears of Dutch-belted rabbits over a frequency range of 0.2 to 16 kHz. Rabbits have a reported 60-dB-SPL hearing range of 0.1 to 50 kHz and most sensitive range of 1 to 16 kHz (Heffner and Masterton, 1980). Therefore, our measurements covered all but the lowest octave and highest 1.6 octaves of rabbit hearing, and covered all of the most sensitive range.

Magnitude, ILD, and ITD spectra of all rabbits are available on a free and public data repository (Day, 2023). Magnitude spectra of source locations in the front horizontal plane had several prominent, stereotypical features (Fig. 3), including: 1) a peak at around 2.6 kHz clearly visible for all azimuths; 2) a sloping portion between approximately 4 and 5 kHz with slope that became more negative as azimuth progressed from 90° ipsilateral (I) to 0°; 3) a notch at around 7.6 kHz visible for most azimuths; and 4) a peak at around 9.8 kHz visible for most azimuths. Before assessing the variability of these spectral features, we first examine the accuracy of measurement of magnitude spectra.

**Fig. 3.**
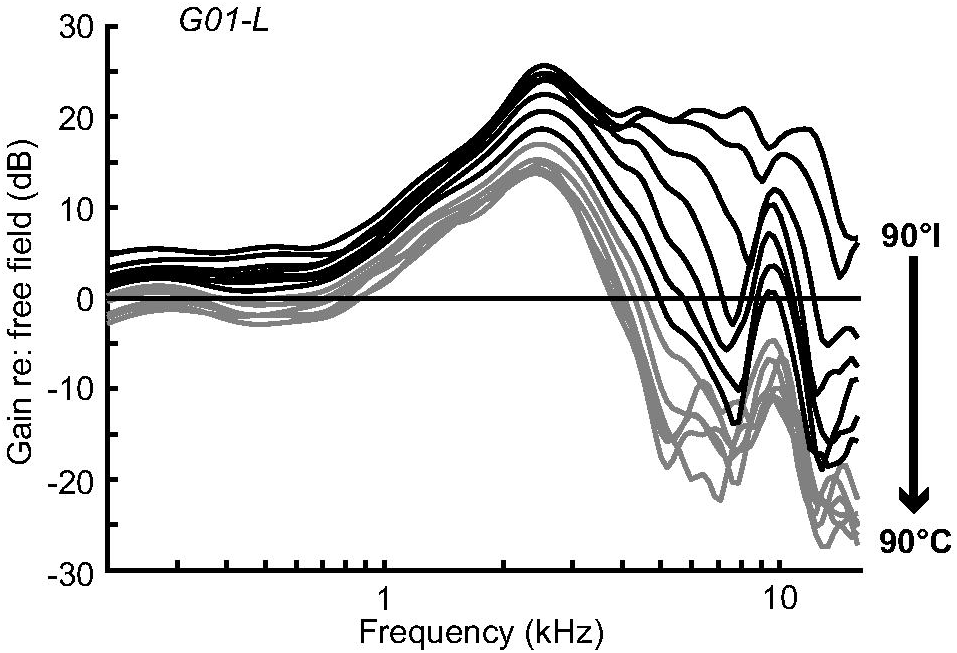
Representative HRTF magnitude spectra from a single rabbit ear, shown for ipsilateral (black; 90°I – 0°) and contralateral azimuths (gray; 15°C – 90°C). Rabbit ID and ear are indicated in the upper-left corner.

### 3.2. Accuracy of magnitude spectra

Figure 4A shows magnitude spectra from the same ear measured twice. Between measurements, we removed the probe tubes from the ears, picked up the rabbit and placed it back down in the correct position, and reinserted the probe tubes into the ears. In this example, test-retest errors in gain were very small below the mid-frequency peak and larger above. We measured HRTFs twice in the same manner in both ears of two rabbits (Fig. 4B). Like the example in Figure 4A, RMS errors across ears were very small below the mid-frequency peak (0.3 dB, on average) and larger above (2.3 dB). One ear showed the largest errors (F04-L), which occurred around 4.5 kHz and 13 kHz; this ear’s magnitude spectra were the examples shown in Figure 4A. The shape of spectra between measurements was very similar for this ear despite the larger error. Altogether, we interpreted measurement error as being relatively small.

**Fig. 4.**
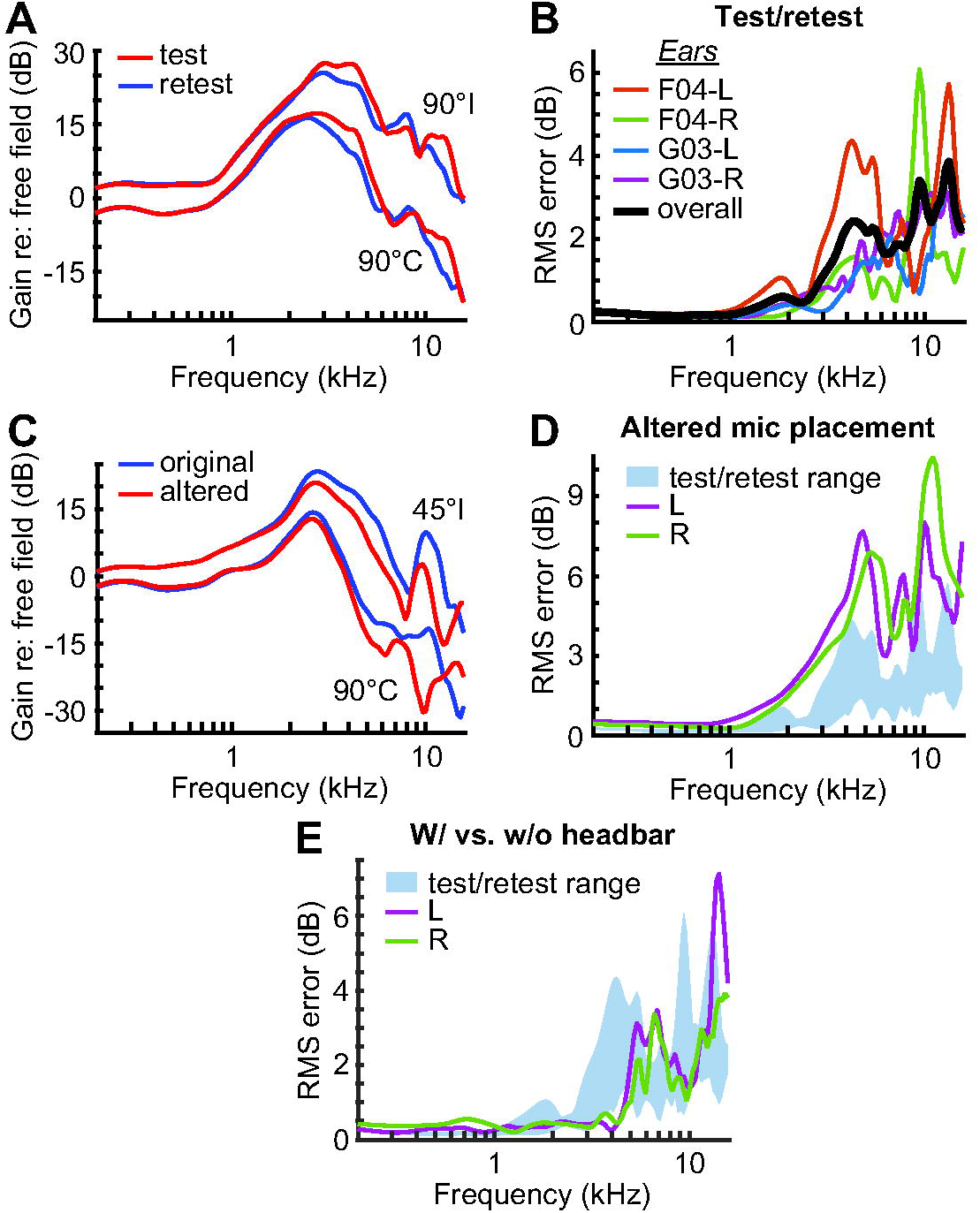
Accuracy of magnitude spectra. (A) Magnitude spectra measured in the same ear twice (test [red] and retest [blue]), shown for two representative source azimuths: 90°I and 90°C. (B) Error between test and retest measurements of magnitude spectra for each ear (different-colored thin lines) and over all ears (black thick line). RMS error was computed at each frequency across source azimuths. (C) Magnitude spectra measured in the same ear using original (blue) and altered (red) microphone positions, shown for two representative source azimuths: 45°I and 90°C. (D) Error between measurements of magnitude spectra using the two microphone positions for the left (purple) and right (green) ears. Range of test/retest errors in (B) shown for comparison. (E) Error between magnitude spectra measured before and after headbar implantation for the left (purple) and right (green) ears.

In one rabbit, we remeasured HRTFs after tugging on the probe tubes to pull them out approximately 5 mm, resulting in a looser fit of the attached shell within the ear. Example magnitude spectra (Fig. 4C) show that spectral shape was mostly the same following this altered microphone placement but could lead to clear differences in some spectral features, such as the conversion of a peak to a notch at 10 kHz for 90° contralateral (C). Altered microphone placement generally led to larger errors above the mid-frequency peak (4.4 dB) than test-retest errors (2.3 dB) (Fig. 4D). Therefore, a snug fit of the shell and attached probe tube within the concha of the rabbit’s ear was important for accurate HRTF measurement.

In another rabbit, we measured HRTFs before and after implantation of a headbar, with five weeks passing between measurements. RMS errors between measurements made before and after headbar implantation overlapped with test-retest measurement error (Fig. 4E). Therefore, we interpreted the effect of a headbar on HRTFs as being insignificant and did not differentiate between data from rabbits with or without headbars in subsequent analyses.

### 3.3. Variability of spectral features across spatial locations and ears

We found the portion of the magnitude spectrum at frequencies lower than the mid-frequency peak had no distinguishing spectral features. Plotting gain relative to that at 0.5 kHz revealed that the shapes of spectra for different azimuths were nearly identical below about 1.5 kHz (Fig. 5A); the SD of relative gain below 1.5 kHz after subtracting the mean at each frequency ranged from only 0.4 to 0.7 dB across ears. Absolute gain at 0.5 kHz increased monotonically from contralateral to ipsilateral azimuths and was similar across ears (Fig. 5B); its SD after subtracting the mean at each azimuth was 0.9 dB. Most of this cross-ear variability was likely due to real, but small, differences across ears because measurement error was only 0.2 dB at 0.5 kHz (Fig. 4B). Putting the analyses together, we found that a change in azimuth simply changed the gain of the magnitude spectrum below 1.5 kHz without changing its shape, and the low-frequency portion of the spectrum was similar across ears.

**Fig. 5.**
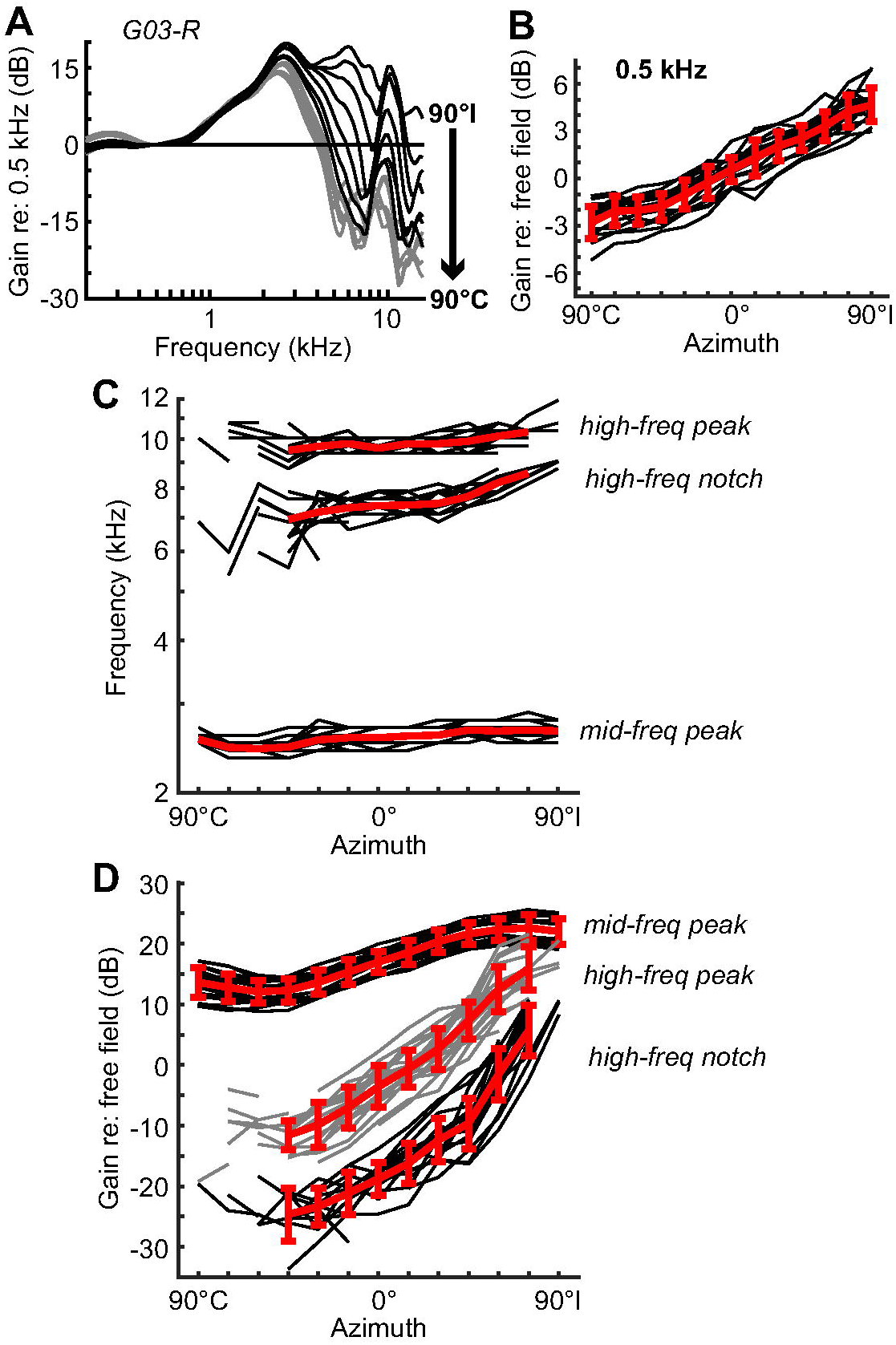
Variation of spectral features with source azimuth. (A) Representative magnitude spectra from a single rabbit ear, each plotted with respect to the gain of the function at 0.5 kHz, shown for ipsilateral (black; 90°I – 0°) and contralateral (gray; 15°C – 90°C) azimuths. (B) Gain of the magnitude spectrum at 0.5 kHz vs. source azimuth; data are shown for each ear (black lines) and the mean ± SD across ears (red). (C, D) Frequencies (C) and gains (D) of the mid-frequency peak, high-frequency notch, and high-frequency peak of the magnitude spectrum vs. source azimuth. Red lines in (C) are the geometric mean frequencies across ears.

Above 1.5 kHz, spectral shape clearly changed for different azimuths (Fig. 5A). Mid-frequency peaks were clearly visible in all ears at all azimuths, whereas high-frequency notches and peaks were clearly visible in most ears only between 45°C and 75°I. The frequencies of mid-frequency and high-frequency peaks were constant across azimuths, whereas the frequency of the high-frequency notch slightly increased at the most ipsilateral azimuths (Fig. 5C). Peak and notch frequencies were highly consistent across ears (Table 1). The gain of the mid-frequency peak varied approximately sinusoidally with azimuth (Fig. 5D). Gains of the high-frequency notch and peak increased from contralateral to ipsilateral azimuths at similar rates such that the peak-to-notch ratio was similar between most azimuths. Cross-ear variability of gain increased for different points of interest along the magnitude spectrum, from the gain at 0.5 kHz (SD = 0.9 dB, described above) to the mid-frequency peak to the high-frequency notch and peak (Table 1). At all points except the high-frequency peak, cross-ear variability of gain was greater than measurement error at the same frequency (Fig. 4B), meaning that at least some of the variability was likely due to real differences across ears.

**Table 1.** Cross-ear variability of spectral features.

|  | **Geometric mean**<br>**frequency (kHz)** | **Geometric SD of**<br>**frequency (oct)** | **SD of gain (dB)** |
| --- | --- | --- | --- |
| *Mid-frequency peak* | 2.6 | 0.05 | 1.9 |
| *High-frequency notch* | 7.6 | 0.08 | 3.5 |
| *High-frequency peak* | 9.8 | 0.05 | 3.2 |

SD's were computed after subtracting the mean at each azimuth.

### 3.4. Utility of monaural magnitude spectra for localization within the front horizontal plane

One consequence of the low-frequency portion of the magnitude spectrum being the same shape across azimuths is that rabbits should not be able to localize sounds within the front horizontal plane using one ear if the sources only contain frequencies below 1.5 kHz. For such sounds, the gain would change with azimuth, but a change in gain could not be disambiguated from a change in intensity of the source of the sound. However, the shape of the magnitude spectrum *above* 1.5 kHz clearly changes with azimuth (Fig. 5A). Theoretically, it should be possible for rabbits to localize sounds within the front horizontal plane using one ear for flat-spectrum sources containing a broad band of frequencies above 1.5 kHz, so long as the shape of the magnitude spectrum at one azimuth is distinct from those at other azimuths (Neti et al., 1992). Determining changes in spectral shape would require the rabbit’s brain to make a comparison of acoustic power between two or more frequency bands (Zakarauskas and Cynader, 1993). We chose to analyze monaural spectral cues for azimuth via a simple difference in the average gain of neighboring frequency bands. This choice was based on previous observations that 1) neurons in central auditory brain areas are topographically organized by characteristic frequency, and 2) many central auditory neurons show frequency tuning that contains neighboring excitatory and inhibitory subfields (Ramachandran et al., 1999), likely due to interactions between neurons in neighboring tonotopic regions. The activity of neurons with excitatory and inhibitory frequency subregions should provide information on spectral contrast and, potentially, sound source location (Imig et al., 1997).

Figure 6A illustrates our division of the upper frequency range into equal-width bands, chosen so that the high-frequency notch and peak fell into the last two bands. For the two comparisons that included the mid-frequency peak, “*i* – *ii*” and “*ii* – *iii*”, the difference of average gains was mostly a monotonic function of azimuth, but the unambiguous portions of the curves only spanned 2 dB and 3 dB, respectively (Fig. 6B and C). The comparison along the slope of the magnitude spectrum between 3 and 5 kHz, “*iii* – *iv*”, had a function that was constant across contralateral azimuths, but decreased monotonically over ipsilateral azimuths by 10 dB (Fig. 6D). The last three comparisons, “*iv* – *v*”, “*v* – *vi*”, and “*vi* – *vii*”, had functions that were nonmonotonic over most azimuths, meaning most values of the difference of average gains could be associated with two or more azimuths (Fig. 6E–G). The comparison “*iv* – *v*” had a small portion of the function between 60°I and 90°I that decreased monotonically, but only spanned 4 dB. Altogether, only the difference of average gains between neighboring bands falling along the spectrum slope between 3 and 5 kHz provided a sizeable, unambiguous cue for a large range of azimuths, limited to the ipsilateral side. The cross-ear variability of the difference of average gains in this comparison over ipsilateral azimuths was relatively small; the SD after subtracting the mean at each azimuth was 0.7 dB.

**Fig. 6.**
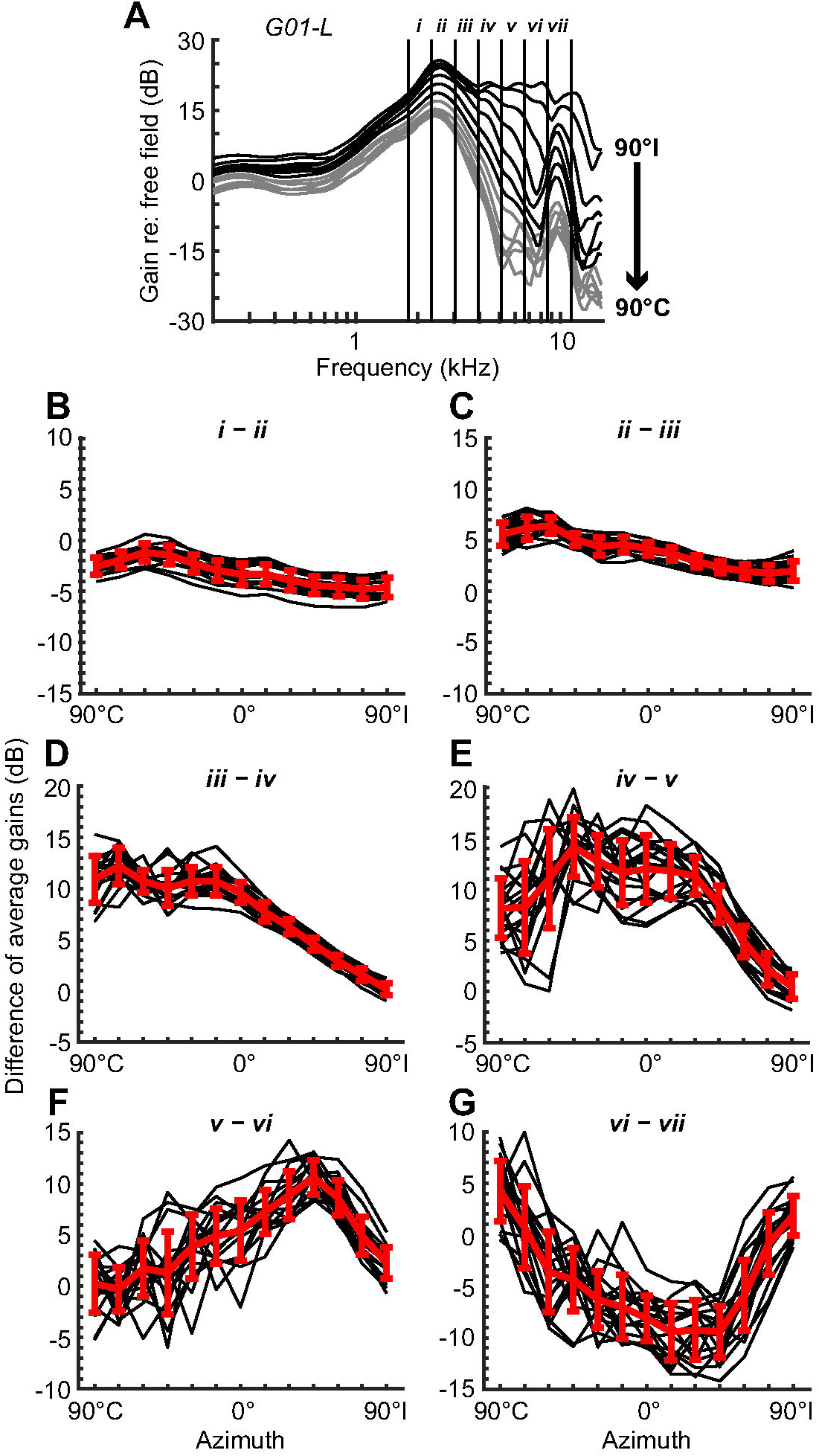
Variation of monaural spectral contrast with source azimuth. (A) Representative magnitude spectra from a single rabbit ear showing the division of the upper frequency range into seven bands of equal width (*i–vii*). (B–G) Difference between the average gains of neighboring frequency bands vs. source azimuth; data are shown for each ear (black lines) and the mean ± SD across ears (red lines). Frequency bands are indicated at the top of each panel.

### 3.5. Variability of binaural cues across spatial locations and ears

We found that ILD spectra for locations within the front horizontal plane could be divided into three frequency regions with characteristic spectral shapes (Fig. 7A). First, a low-frequency region below about 1.5 kHz with a flat spectrum whose value changed with azimuth. This region corresponded to the frequency region of magnitude spectra where spectral shape remained constant across azimuths. Second, a region between about 1.5 and 5 kHz where ILD magnitude increased smoothly with frequency. This region overlapped with the mid-frequency peak and spectral slopes of magnitude spectra. Third, a high-frequency region above about 5 kHz where ILD spectra took on complex shapes that were inconsistent across azimuths. This last region overlapped with the high-frequency notch and peak of magnitude spectra. Although the high-frequency shapes of ILD spectra were dissimilar across azimuths, they did show some similarity for the same azimuth across ears (Fig. 7C).

**Fig. 7.**
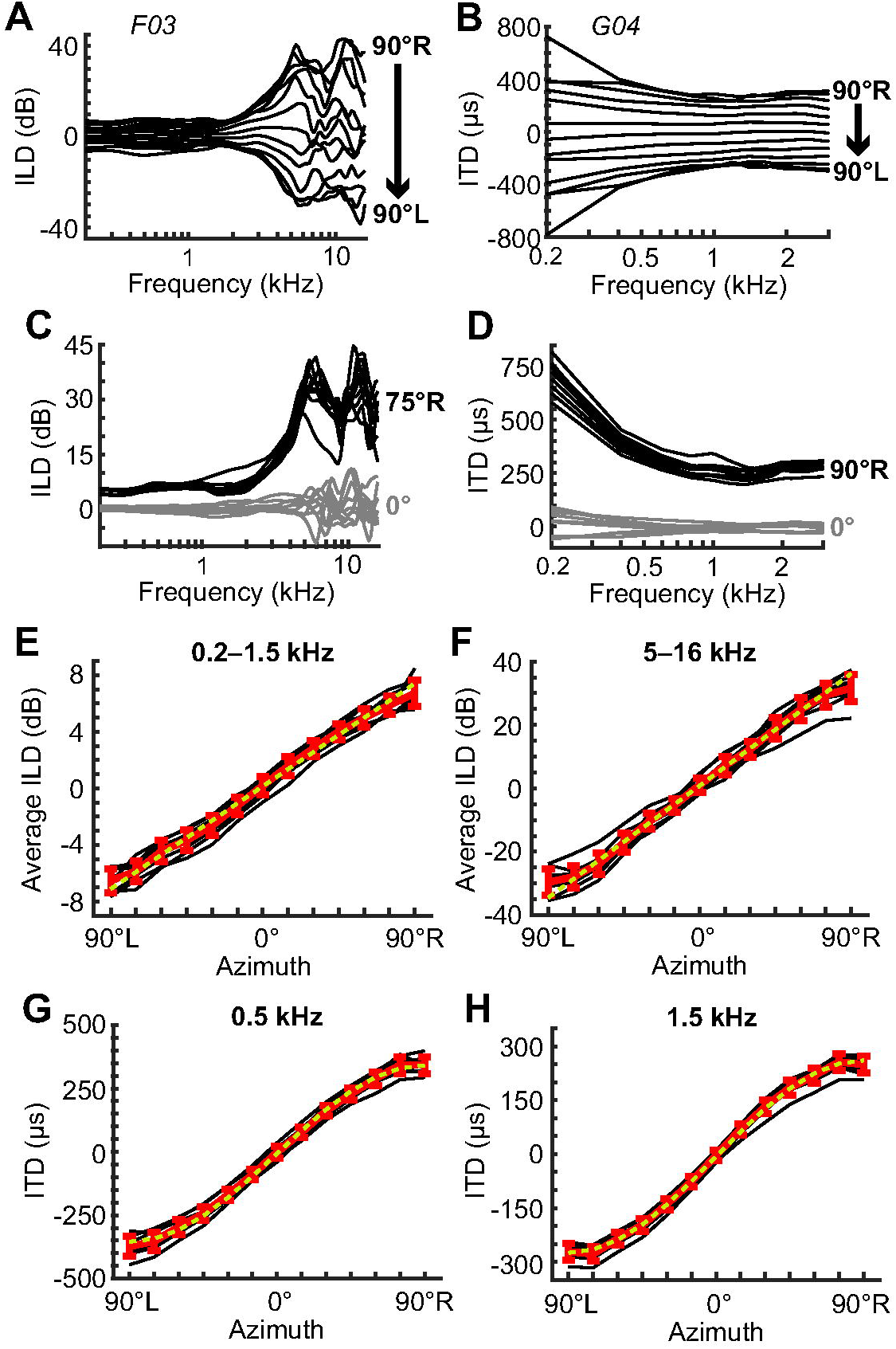
Variation of binaural cues with source azimuth. (A, B) Representative ILD (A) and ITD spectra (B) from single rabbits, shown for azimuths within the front horizontal plane. (C, D) ILD (C) and ITD spectra (D) of all rabbits for sources at 0° (gray) and 75°R or 90°R (black; C and D, respectively). (E, F) Average ILD vs. source azimuth computed within low-frequency (E; 0.2–1.5 kHz) or high-frequency bands (F; 5–16 kHz); data are shown for individual rabbits (black lines), the mean ± SD across rabbits (red lines), and fit lines (yellow dashed lines). (G, H) ITD vs. source azimuth at 0.5 kHz (G) and at 1.5 kHz (H); yellow dashed lines are sinusoidal fits.

We computed the functions of average ILD vs. azimuth for a low-frequency band (0.2– 1.5 kHz; Fig. 7E) and a high-frequency band (5–16 kHz; Fig. 7F). Unlike monaural spectral cues, average ILD was monotonic and increasing over the whole range of azimuths, forming a one-to-one mapping of ILD to azimuth. Functions were well fit by a line except for slight compression at 90°L and R. The absolute value of ILD at 90°L and R was 6.6 ± 0.9 dB (mean ± SD) for the low-frequency band and 31 ± 4 dB for the high-frequency band. In comparison, the test-retest error for the same azimuths were 0.4 dB and 3.6 dB for the low- and high-frequency bands, respectively. Therefore, most of the cross-ear variability in ILD was likely due to real differences between rabbits for the low-frequency band, but not the high-frequency band.

We found that ITD spectra for locations within the front horizontal plane could be characterized into two frequency regions (Fig. 7B): 1) a region below about 1 kHz where ITD magnitude increased with decreasing frequency, and 2) a region above about 1 kHz where spectra were flat. The smooth, monotonic shapes of ITD spectra were simple and highly consistent across ears (Fig. 7D). Functions of ITD vs. azimuth at 0.5 kHz and 1.5 kHz were sinusoidally shaped (Fig. 7G and H); like ILD, there was a one-to-one mapping of ITD to azimuth. The absolute value of ITD at 90°L and R was 358 ± 38 μs (mean ± SD) at 0.5 kHz and 260 ± 25 μs at 1.5 kHz. In comparison, the test-retest error for the same azimuths was 9 μs and 13 μs for 0.5 kHz and 1.5 kHz, respectively. Therefore, most of the cross-ear variability in ITD was likely due to real differences between rabbits.

### 3.6. Contribution of pinnae to acoustic filtering

In order to quantify the contribution of the pinnae to acoustic filtering, we measured HRTFs in a cadaver rabbit before and after removal of its pinnae. The resulting magnitude spectra following pinna removal lacked all of the characteristic features of magnitude spectra; they had no mid-frequency peak, no mid-frequency sloping portion, and no high-frequency notch or peak (Fig. 8A). Magnitude spectra following pinna removal did show gain at middle frequencies and attenuation at high frequencies for contralateral locations, but not as large in amplitude as magnitude spectra from intact pinnae. The gain at middle frequencies in magnitude spectra lacking pinnae was only 32% of the gain of the mid-frequency peak in magnitude spectra from intact pinnae, on average. At low frequencies, magnitude spectra from intact pinnae were slightly higher than magnitude spectra lacking pinnae for many ipsilateral azimuths (Fig. 8A), indicating a small contribution of the pinnae even at low frequencies.

**Fig. 8.**
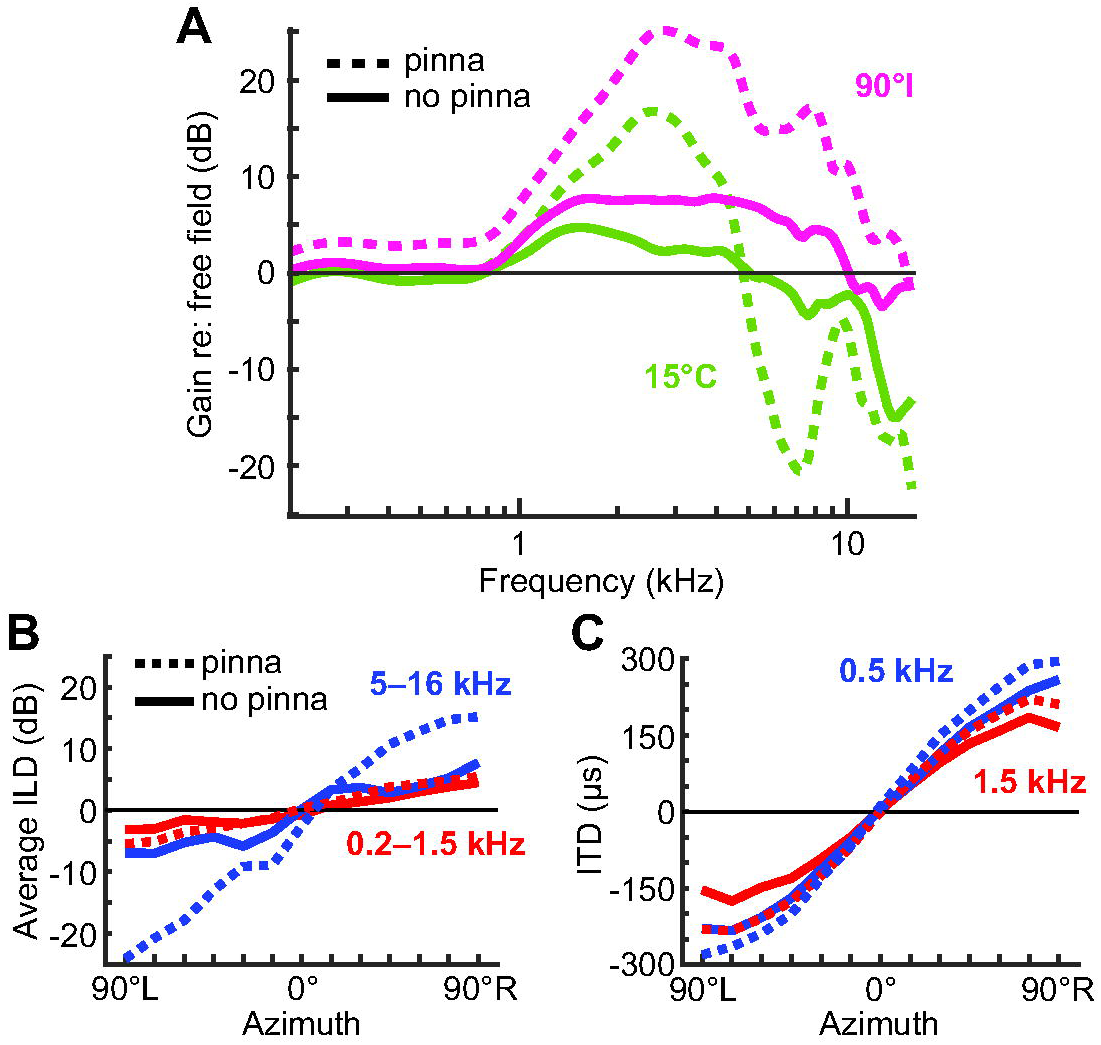
Effect of pinna removal. (A) Magnitude spectra of one ear of a cadaver rabbit for sources at 90°I (magenta) and 15°C (green), measured before (dashed lines) and after (solid line) removal of pinnae. (B, C) Average ILD (B) and ITD (C) vs. source azimuth measured before (dashed lines) and after (solid lines) removal of pinnae. (B) Average ILD was computed within frequency bands of 0.2–1.5 kHz (red) and 5–16 kHz (blue). (C) ITD is shown for frequencies of 0.5 kHz (blue) and 1.5 kHz (red).

ILD in the low-frequency band (0.2–1.5 kHz) was less affected by pinna removal than ILD in the high-frequency band (5–16 kHz) (Fig. 8B). For the low-frequency band, the absolute value of ILD averaged between 90°L and R was 5.5 dB with pinnae and 3.8 dB without pinnae (or 70% of the value for intact pinnae). For the high-frequency band, the absolute value of ILD averaged between 90°L and R was 19.6 dB with pinnae and 7.3 dB without pinnae (or 37% of the value for intact pinnae). ITD was only weakly affected by pinna removal (Fig. 8C). The absolute value of ITD averaged between 90°L and R was 288 μs with pinnae and 244 μs without pinnae (or 85% of the value for intact pinnae) at 0.5 kHz, and 220 μs with pinnae and 160 μs without pinnae (or 72% of the value for intact pinnae) at 1.5 kHz. Altogether, our data from the cadaver rabbit demonstrate that pinnae create the characteristic spectral features of magnitude spectra and are the largest contribution to high-frequency ILD, whereas the rest of the head is the largest contribution to ITD and low-frequency ILD.

### 3.7. Dependence of acoustic filtering on sex, mass, or age

We did not make anatomical measurements of the head or pinnae of rabbits; however, we did record the sex, mass, and age of rabbits in the study, which may be factors associated with head and pinna size. We tested for possible effects of these factors on three measured quantities for which cross-ear variability was much greater than measurement error: the gain at 0.5 kHz for 90°I, the gain of the mid-frequency peak for 90°I, and the absolute value of ITD at 1.5 kHz for 90°L and R. For all quantities, values from each individual rabbit were averaged between left and right ears. We found no sex differences for any of the three quantities (Table 2); however, sample size was small, reducing statistical power. The gain of the mid-frequency peak was positively correlated with mass, and the gain at 0.5 kHz was negatively correlated with age; otherwise, there were no other significant correlations with mass or age.

**Table 2.**
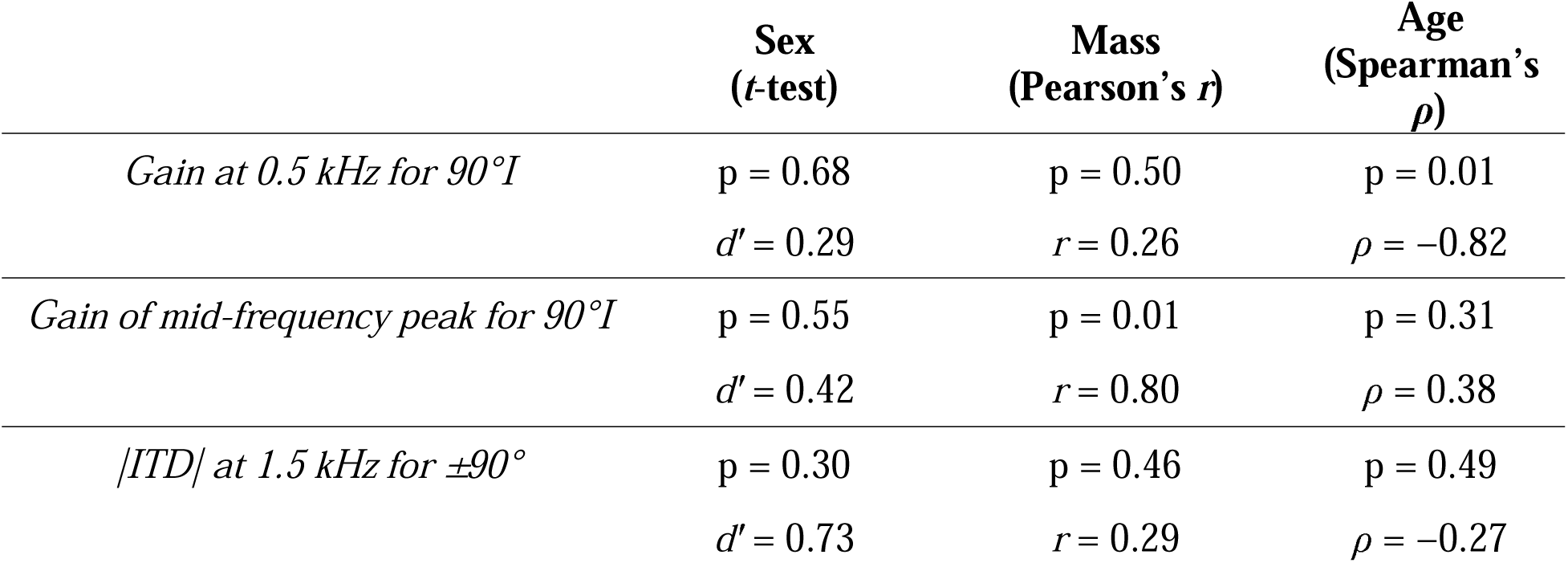
Effect of sex, mass, or age on HRTF quantities.

### 3.8. Similarity of magnitude spectra between rabbits

Figure 9A shows the magnitude spectra of all ears in our sample for three source locations; the shapes of magnitude spectra were qualitatively similar across individual ears. To quantify the overall similarity of magnitude spectra between ears of different rabbits, we took each possible pair of matched-location magnitude spectra between ears of different rabbits and computed the RMS error over all azimuths and frequencies for the pair. We quantified the similarity of magnitude spectra between ears of the same rabbit in the same manner. The RMS error of different-rabbit pairs of ears was significantly greater than the RMS error of same-rabbit pairs of ears (Fig. 9B; p = 0.002, *t*-test), indicating that, on average, magnitude spectra of a rabbit’s ear were more similar to those of its other ear than to those of other rabbits’ ears.

**Fig. 9.**
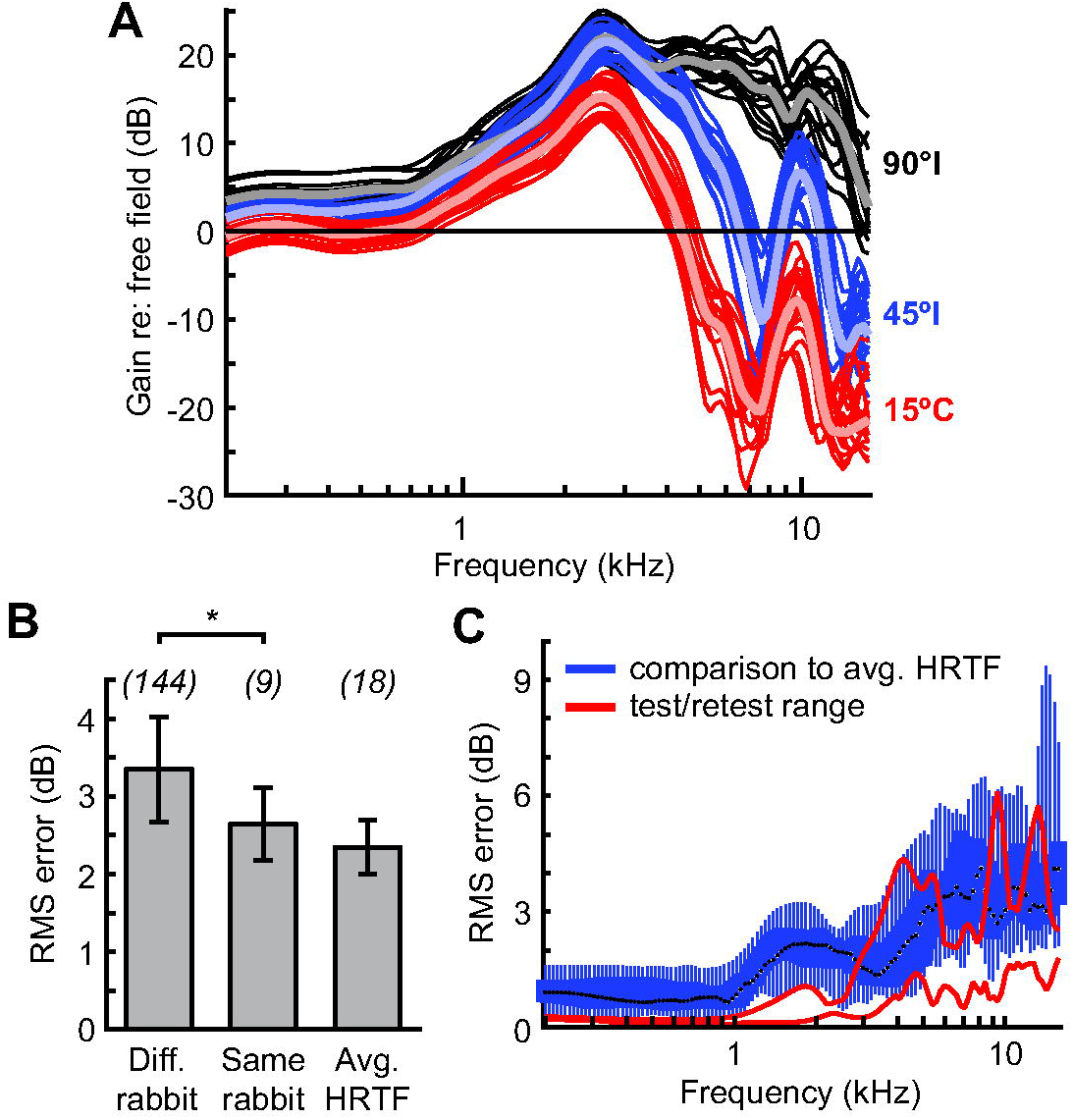
Similarity of magnitude spectra. (A) Magnitude spectra of all rabbit ears (thin lines) and the average rabbit ear (thick lines) for sources at 90°I (black), 45°I (blue), and 15°C (red). (B) RMS error (mean ± SD) between magnitude spectra of different-rabbit pairs of ears, same-rabbit pairs of ears, and individual rabbit ears and the average rabbit ear. Error was computed across frequencies and azimuths for each pair. Numbers in parentheses indicate the number of pairs. Asterisk indicates p < 0.05 (*t*-test). (C) RMS error as a function of frequency between magnitude spectra of individual rabbit ears and the average rabbit ear, computed across azimuths for each pair. At each frequency is a box plot showing median (black circles), interquartile range (thick blue lines), and range (thin blue lines). Range of test/retest errors (red lines) from Fig. 4B shown for comparison.

We also quantified the similarity of magnitude spectra between individual ears and an average ear. RMS error was not significantly different than that for same-rabbit pairs of ears (Fig. 9B; p = 0.08, *t*-test), indicating that, on average, magnitude spectra of a rabbit’s ear were as similar to those of an average ear as to those of its other ear. The RMS error between magnitude spectra of individual ears and an average ear increased with frequency but remained relatively small (Fig. 9C). RMS error overlapped with the range of measurement error in several frequency subregions above 3 kHz. We confirmed that quantitative values of the properties of average HRTFs were identical to or nearly identical to the mean values of these properties across ears, including: the gain at 0.5 kHz as a function of azimuth (Fig. 5B); the peak frequency and gain of the mid-frequency peak, high-frequency notch, and high-frequency peak as a function of azimuth (Fig. 5C and D); low-frequency and high-frequency ILD as a function of azimuth (Fig. 7E and F); and ITD at 0.5 kHz and 1.5 kHz as a function of azimuth (Fig. 7G and H). In other words, the standard deviations in plots referenced in the preceding sentence, which are deviations from the mean, are also the same as the standard deviations of cross-ear data from the average HRTF.

## 4. Discussion

### 4.1. Validity of HRTFs

The HRTF is the pressure transformation from free-field space to tympanic membrane; however, measurements are not of pressure at the tympanic membrane itself but of air pressure at a point near the tympanic membrane. This raises the possibility that HRTFs may not accurately capture the pressure at the input to the auditory system. Previous studies have discussed this issue in detail (see Discussion of Rice et al., 1992). In humans, measurements from two probe microphones placed within the ear canal a fixed distance away from each other were found to have magnitude spectra that were different in *absolute* terms but the same in their *directional dependence* (Middlebrooks et al., 1989). In other words, the ratio of magnitude spectra between two different spatial locations was the same between the two probe microphone positions. This result was consistent with acoustic modeling in humans and cats that showed pressure waves that successfully traverse the extent of the ear canal are in the form of plane waves (i.e., equal pressure over the cross-section of the canal; Rabbitt and Holmes, 1988; Rice et al., 1992). Therefore, absolute pressure may differ across positions within the ear canal due to standing wave patterns and complex modes of vibration near the tympanic membrane, but the dependence of pressure on sound source location is the same across positions within the ear canal since directional filtering occurs outside of the ear canal. Given that the frequencies used in the present study (<16 kHz) were much less than lower frequency limits at which directional dependence of pressure transformations due to non-plane-wave propagation are predicted to occur in humans and cats (28 kHz and 42 kHz, respectively; Rabbitt and Holmes, 1988; Rice et al., 1992), it is likely that the directional dependence of our data in rabbits is accurate.

Since the pressure measured by a probe microphone in the ear canal may not be the same as that at the tympanic membrane in absolute terms, researchers often report the directional transfer function (DTF), which is the ratio of an HRTF magnitude spectrum to the average magnitude spectrum across all spatial locations (Koka et al., 2008; Maki and Furukawa, 2005; Middlebrooks and Green, 1990). We did not compute DTFs because we only sampled spatial locations within the front horizontal plane, which would have led to a skewed average magnitude spectrum. Alternatively, we computed the ratio of magnitude spectra at various azimuths to that at a reference azimuth of 0°, which eliminates the potentially erroneous nondirectional component of the HRTF; two of such relative magnitude spectra are shown in Figure 10A (average HRTFs). The high-frequency notch and peak in HRTF magnitude spectra shifted to slightly higher frequencies in the relative magnitude spectra, and an additional prominent peak appeared below the notch, all of which leads to a confusing interpretation. Relative magnitude spectra lack some scientifically important information, such as the ear canal resonance and whether the pinna amplifies or attenuates the signal with respect to free field at specific frequencies and source azimuths. Therefore, we chose to report absolute HRTF data.

**Fig. 10.**
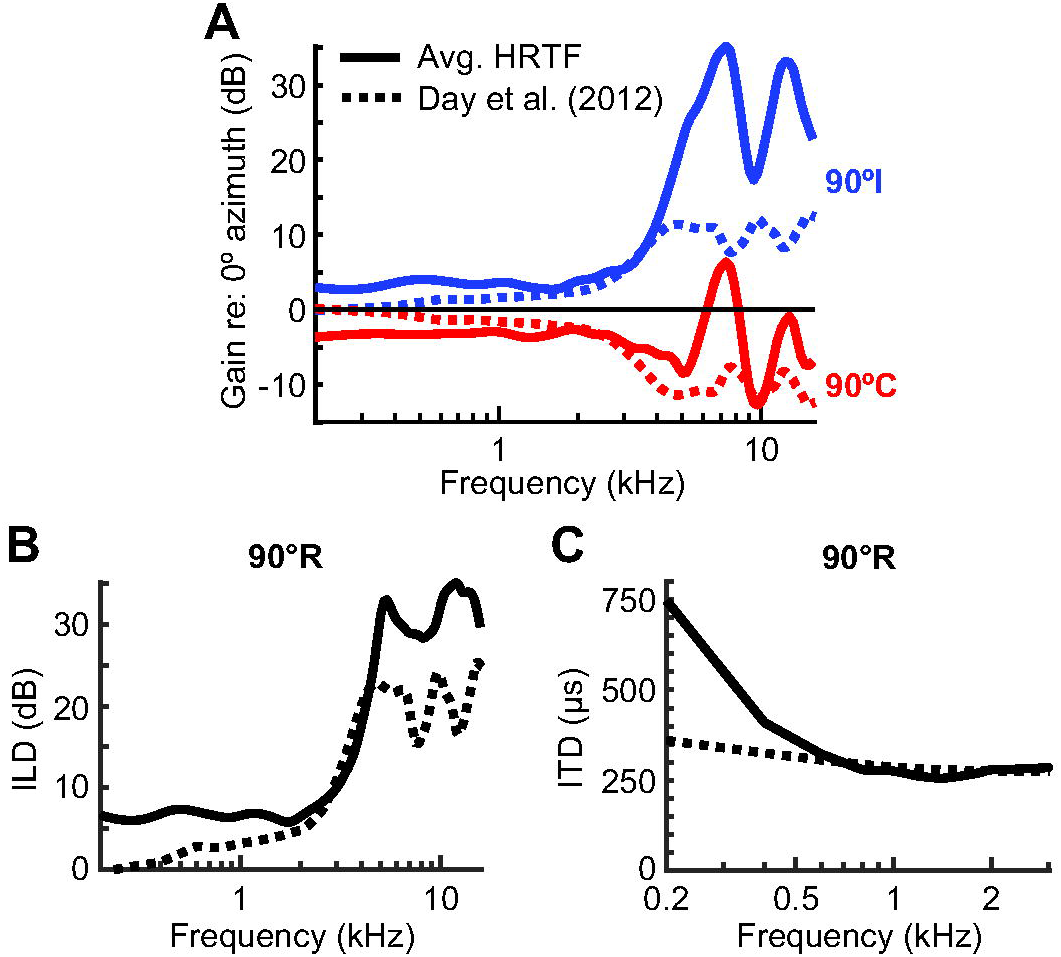
Comparison to previous non-individualized DTFs for rabbits. (A) The ratio of magnitude spectra between source azimuths of 90°I and 0° (blue) and the ratio of magnitude spectra between source azimuths of 90°C and 0° (red). Taking the ratio between magnitude spectra of specific locations and a reference location (0°) removes the nondirectional components of HRTFs, allowing comparison with DTFs. Data are shown for the average HRTF in the present study (solid lines) and the DTFs computed in a previous study (dashed lines; Day et al., 2012). (B, C) ILD (B) and ITD (C) spectra for a source azimuth of 90°R.

We found that tugging on the probe microphone tube to move the position of its tip within the ear by approximately 5 mm changed magnitude spectra but left characteristics of prominent spectral features largely unaltered, including the frequency of the ear canal resonance, the direction-dependent slope between 3 and 5 kHz, the frequency of the high-frequency notch and peak, and the overall shift in high-frequency gain from amplification to attenuation with changes in source azimuth from ipsilateral to contralateral. That these spectral features remained after a relatively large shift in probe microphone position increases confidence that the same spectral features are present in the input to the auditory system. Previous studies have shown a prominent, nondirectional notch in the magnitude spectrum that shifts in frequency with probe microphone position, likely due to a standing wave null within the ear canal (Maki and Furukawa, 2005; Shaw and Teranishi, 1968). Our data showed no such nondirectional notch, which suggests that the standing wave null for our probe microphone placement, if it existed, was above our highest measured frequency.

### 4.2. Comparison of spectral features of magnitude spectra to those of other species

We found a broad spectral peak around 2.6 kHz whose peak frequency and shape was unaffected by source location, but whose gain increased from 12 dB for contralateral azimuths to 23 dB for ipsilateral azimuths, on average, thereby amplifying a large range of frequencies surrounding the peak for all source locations. The same peak was reported for one rabbit in a previous study (Kim et al., 2010). Similar broad spectral peaks have been observed in many other mammals and interpreted as an acoustic resonance along the length of the ear canal (Shaw and Teranishi, 1968). The absence of the peak following pinna removal (Fig. 8A), which left only the bony portion of the ear canal, demonstrates that the resonance in rabbits resulted from the pinna, including the concha and cartilaginous portion of the ear canal. Peak frequency of the ear canal resonance in rabbits was similar to that of humans (2–3 kHz; Shaw, 1974) and chinchillas (∼2.7 kHz; Koka et al., 2011), but less than that of cats (3–5 kHz; Rice et al., 1992), guinea pigs (∼5.5 kHz; Greene et al., 2014), marmosets (6–12 kHz; Slee and Young, 2010), gerbils (∼8 kHz; Maki and Furukawa, 2005), ferrets (∼8 kHz; Carlile, 1990b), mice (10–20 kHz; Lauer et al., 2011), and rats (∼18 kHz; Koka et al., 2008). The overlap of the resonant frequencies of rabbits and chinchillas with that of humans, who have a relatively large average ear canal length (26 mm; Goode et al., 1977), may be due to the relatively large pinnae of the smaller-headed mammals.

In rabbits, frequencies above 4 kHz were amplified or attenuated with respect to free field depending on the azimuth of the sound source: frequencies were most amplified at 90°I and most attenuated between 45°C and 90°C (Fig. 3 and 5D). This directionality of gain was mostly due to the pinna (Fig. 8A) and was consistent with the concavity of the pinna in resting position facing the 90°I source, assuming the pinna acts as a directional sound collector. The movability of rabbit pinnae therefore allows rabbits the ability to selectively amplify sources in the direction of the pinna concavity. The directionality of high-frequency gain is also apparent in data from one rabbit in a previous study where pinnae were oriented upright with the concavity facing laterally (Kim et al., 2010). Similar directionally dependent, high-frequency gain is observed in other species; the lower frequency bound of directional gain in rabbits (∼4 kHz) was similar to that of humans (5–6 kHz; Shaw, 1974; Wightman and Kistler, 1989) and cats (∼5 kHz; Rice et al., 1992), but less than that of marmosets (∼10 kHz; Slee and Young, 2010).

We observed a prominent spectral notch for most source azimuths at around 7.6 kHz. Similar spectral notches have been observed in other species and named the “first notch” due to it being the notch with the lowest notch frequency. These studies typically report multiple notches in the HRTF magnitude spectrum (Rice et al., 1992; Shaw and Teranishi, 1968; Slee and Young, 2010); it is possible we would have also encountered additional spectral notches had we measured at frequencies above 16 kHz. First-notch frequency took on a range of 6.5 to 9 kHz across rabbits, similar to that of humans (∼8 kHz; Shaw and Teranishi, 1968), slightly less than that of cats (9–16 kHz; Rice et al., 1992), and much less than that of marmosets (12–24 kHz; Slee and Young, 2010). First-notch frequency was also mostly stable across source azimuths, similar to humans (Shaw and Teranishi, 1968) and marmosets (Slee and Young, 2010), but unlike cats (Rice et al., 1992), which have first notches that systematically shift in frequency within the front horizontal plane. Previous studies have uniformly found the first-notch frequency to vary substantially with source elevation (Rice et al., 1992; Shaw and Teranishi, 1968; Slee and Young, 2010).

### 4.3. Comparison of binaural cues to those of other species

Maximal high-frequency ILD in rabbits (31 dB, on average) was similar to that of humans and many of other mammalian species, which fell within a range of 20–30 dB (Carlile, 1990a; Greene et al., 2014; Koka et al., 2011; Maki and Furukawa, 2005; Middlebrooks et al., 1989; Musicant et al., 1990; Slee and Young, 2010). The maximal high-frequency ILD was larger than that reported in a previous study of data from one rabbit (∼22 dB; Kim et al., 2010), perhaps due to individual variation; one individual in our data set had a similar value (Fig. 7F). Like other species, rabbits exhibited frequency-dependent ILD: ILD increased smoothly with frequency up to a high-frequency region whereby it ceased to vary smoothly due to spectral peaks and notches in the underlying magnitude spectra. The lower-frequency limit of this high-frequency region was around 5 kHz in rabbits, but varies across species and is, notably, approximately the same as that in humans (∼6 kHz; Shaw, 1974). The pinnae were the main contribution to high-frequency ILD in rabbits, as has been shown in guinea pigs (Greene et al., 2014) and rats (Koka et al., 2008), but not chinchillas (Koka et al., 2011).

ITD in rabbits was also frequency-dependent, exhibiting larger magnitudes at lower frequencies than at higher frequencies, as has been shown in humans (Kuhn, 1977) and many other mammalian species (Benichoux et al., 2016). Maximal ITD of rabbits at 1.5 kHz was 260 μs, on average, slightly larger than that reported from one rabbit in a previous study (Kim et al., 2010), much smaller than that of humans (∼650 μs; Kuhn, 1977), and similar to those of other mammals with medium head sizes (guinea pig: 275 μs, Greene et al., 2014; chinchilla: 236 μs, Koka et al., 2011; cat: ∼300 μs, Roth et al., 1980). The pinnae contributed some to ITD in rabbits, but were not the main contribution, as has also been shown in chinchillas (Koka et al., 2011), rats (Koka et al., 2008), and guinea pigs (Greene et al., 2014).

### 4.4. Monaural spectral cues for localization

In humans, monaural spectral cues are known to underlie perception of sound source elevation and to resolve front/back ambiguity in the horizontal plane, whereas ITD and ILD dominate perception of sound source azimuth (Middlebrooks and Green, 1991). The frequency region surrounding the first notch in HRTF magnitude spectra is particularly implicated in vertical localization and discrimination, as has been shown in humans (Langendijk and Bronkhorst, 2002), cats (Huang and May, 1996) and marmosets (Chen et al., 2023). As mentioned above, the frequency of the first notch is known to vary with sound source elevation. We found that in rabbits, the depth, but not the frequency, of the first notch varied with sound source azimuth and led to a systematically decreasing spectral slope at frequencies below the notch, which provided an unambiguous cue for azimuth ipsilateral to the ear. While it is unlikely that rabbits exclusively use this monaural cue, and not ILD, to determine sound source azimuth, spectral contrast along the slope or over the first notch may potentially be utilized in determining elevation. It will be important to repeat our analysis of monaural spectral contrast on HRTF data measured over the whole hemisphere and not simply the front horizontal plane.

### 4.5. Suitability of average HRTFs for use in future studies

HRTF and DTF magnitude spectra in humans (Wightman and Kistler, 1989) and cats (Xu and Middlebrooks, 2000) have been shown to significantly differ in shape across individuals, particularly in the first-notch frequency range. Consistent with inter-individual differences in HRTFs, human subjects listening to sounds in virtual acoustic space make more elevation and front/back errors when using another individual’s HRTFs (Wenzel et al., 1993). Therefore, it is best practice to use an individual’s own HRTFs in studies of sound localization that use virtual acoustic space. However, measurement of individual HRTFs is not feasible for many neurophysiology laboratories, and use of an accurate non-individualized HRTF would be advantageous.

In previous neurophysiological studies, we presented sound to rabbits in virtual acoustic space using non-individualized DTFs (Day and Delgutte, 2013, 2016; Day et al., 2012; Dorkoski et al., 2020; Haragopal et al., 2020). The DTFs were created from data of one cadaver rabbit with pinnae in upright position (Day et al., 2012). Our aim was to create the ability to present sounds with a realistic dependence of ITD and ILD on frequency and azimuth, while avoiding monaural spectral features that may have differed between rabbits. Therefore, we created simplified DTF magnitude spectra based on the first principal component of a principal components analysis of the complete set of DTFs. Human listeners have been shown to determine azimuth using the first principal component as accurately as using their complete HRTFs (Kistler and Wightman, 1992). We created phase spectra by fitting a sinusoid to the function of azimuth vs. ITD measured at 1.5 kHz, then assumed an exponential expansion of ITD at lower frequencies. Figure 10 shows a comparison of binaural and monaural properties between the non-individualized DTFs and average rabbits HRTFs in the present study. The non-individualized DTFs accurately captured the frequency dependence of ILD but underestimated maximal ILD in both the low- and high-frequency ranges. Furthermore, they accurately captured maximal ITD at high frequencies, but slightly underestimated maximal ITD at 0.5 kHz. Lastly, the directional dependence of magnitude spectra was not accurately captured, which was expected from DTFs based solely on the first principal component.

Inter-individual differences in magnitude spectra, ILD, and ITD were all small in our sample of rabbits. Although deviations of individual magnitude spectra from average spectra increased in the high-frequency range containing the first notch, the frequencies of the first notch and high-frequency peak showed little variation across ears. We believe the average HRTFs in the present study for spatial locations in the front horizontal plane are a reasonably accurate substitute for individualized HRTFs in neurophysiological or behavioral experiments on rabbits utilizing virtual acoustic space. To that end, we provide our average HRIRs in an online repository (Day, 2023), which may be used to filter sounds presented to rabbits at the entrance of the ear canal via a closed field, such as sealed ear inserts. Although we did not find consistent evidence for an effect of sex, mass, or age on properties of the HRTF, the range of masses in our sample was small and all rabbits were adult. Properties of the HRTF have been shown to correlate with head size and pinna size and shape (Middlebrooks, 1999; Schnupp et al., 2003). Therefore, to minimize experimental error, the average HRIRs should be used with rabbits similar to those in the present study: adults from the Dutch-belted strain of similar mass.

## 5. Conclusions

- The rabbit pinna creates a resonance that amplifies a broad range of frequencies around 2.6 kHz regardless of source direction, similar to that of humans.
- The pinna also causes directionally dependent gain of frequencies above 4 kHz, amplifying sources ipsilateral to the pinna concavity and attenuating sources contralateral to the pinna concavity.
- The “first notch” in acoustic filtering performed by the pinna occurs at a frequency of around 7.6 kHz, mostly stable across source azimuths, and similar to that of humans.
- The slope of magnitude spectra between 3 and 5 kHz is an unambiguous monaural cue for source azimuths ipsilateral to the ear.
- ILD and ITD are unambiguous cues for source azimuth in the front horizontal plane, with ranges of ±31 dB (high-frequency) and ±358 μs (0.5 kHz), respectively.
- The pinnae are the main contribution to high-frequency ILD, but are a minor contribution to ITD.
- Variability across rabbits of both monaural spectral features and binaural cues is relatively small, suggesting that an average HRTF may be an adequate substitute for individual HRTFs in experiments on rabbits in virtual acoustic space.

## Acknowledgments

We thank Ken Hancock for writing software code for control of stimulus presentation and data acquisition, and Emili Garretson for assistance with experiments. This work was supported by the National Institute on Deafness and Other Communication Disorders of the National Institutes of Health (R15DC017616).

## Notes

https://doi.org/10.6084/m9.figshare.24085452.v2

### Competing Interest Statement

The authors have declared no competing interest.

## References

Barzelay, O., David, S., Delgutte, B., 2023. Effect of Reverberation on Neural Responses to Natural Speech in Rabbit Auditory Midbrain: No Evidence for a Neural Dereverberation Mechanism. eNeuro 10(5). 10.1523/ENEURO.0447-22.2023.

Benichoux, V., Rebillat, M., Brette, R., 2016. On the variation of interaural time differences with frequency. J Acoust Soc Am 139(4), 1810. 10.1121/1.4944638.

Borg, E., Engstrom, B., Linde, G., Marklund, K., 1988. Eighth nerve fiber firing features in normal-hearing rabbits. Hear Res 36(2-3), 191–201.

Carlile, S., 1990a. The auditory periphery of the ferret. I: Directional response properties and the pattern of interaural level differences. J Acoust Soc Am 88(5), 2180–2195. 10.1121/1.400115.

Carlile, S., 1990b. The auditory periphery of the ferret. II: The spectral transformations of the external ear and their implications for sound localization. J Acoust Soc Am 88(5), 2196–2204. 10.1121/1.400116.

Carney, L.H., Zilany, M.S., Huang, N.J., Abrams, K.S., Idrobo, F., 2014. Suboptimal use of neural information in a mammalian auditory system. J Neurosci 34(4), 1306–1313. 10.1523/JNEUROSCI.3031-13.2014.

Chen, C., Remington, E.D., Wang, X., 2023. Sound localization acuity of the common marmoset (Callithrix jacchus). Hear Res 430, 108722. 10.1016/j.heares.2023.108722. [dataset]

Day, M., 2023. Rabbit head-related transfer functions within the front horizontal plane. 10.6084/m9.figshare.24085452.v2.

Day, M.L., Delgutte, B., 2013. Decoding sound source location and separation using neural population activity patterns. J Neurosci 33(40), 15837–15847. 10.1523/JNEUROSCI.2034-13.2013.

Day, M.L., Delgutte, B., 2016. Neural population encoding and decoding of sound source location across sound level in the rabbit inferior colliculus. J Neurophysiol 115(1), 193–207. 10.1152/jn.00643.2015.

Day, M.L., Koka, K., Delgutte, B., 2012. Neural encoding of sound source location in the presence of a concurrent, spatially separated source. J Neurophysiol 108(9), 2612–2628. 10.1152/jn.00303.2012.

Dorkoski, R., Hancock, K.E., Whaley, G.A., Wohl, T.R., Stroud, N.C., Day, M.L., 2020. Stimulus-frequency-dependent dominance of sound localization cues across the cochleotopic map of the inferior colliculus. J Neurophysiol 123(5), 1791–1807. 10.1152/jn.00713.2019.

Fan, L., Henry, K.S., Carney, L.H., 2022. Responses to dichotic tone-in-noise stimuli in the inferior colliculus. Front Neurosci 16, 997656. 10.3389/fnins.2022.997656.

Goode, R.L., Friedrichs, R., Falk, S., 1977. Effect on hearing thresholds of surgical modification of the external ear. Ann Otol Rhinol Laryngol 86(4 Pt 1), 441–450. 10.1177/000348947708600404.

Greene, N.T., Anbuhl, K.L., Williams, W., Tollin, D.J., 2014. The acoustical cues to sound location in the guinea pig (Cavia porcellus). Hear Res 316, 1–15. 10.1016/j.heares.2014.07.004.

Haragopal, H., Dorkoski, R., Pollard, A.R., Whaley, G.A., Wohl, T.R., Stroud, N.C., Day, M.L., 2020. Specific loss of neural sensitivity to interaural time difference of unmodulated noise stimuli following noise-induced hearing loss. J Neurophysiol 124(4), 1165–1182. 10.1152/jn.00349.2020.

Heffner, H., Masterton, B., 1980. Hearing in Glires: Domestic rabbit, cotton rat, feral house mouse, and kangaroo rat. J Acoust Soc Am 68, 1584–1599.

Huang, A.Y., May, B.J., 1996. Sound orientation behavior in cats. II. Mid-frequency spectral cues for sound localization. J Acoust Soc Am 100(2 Pt 1), 1070–1080. 10.1121/1.416293.

Imig, T.J., Poirier, P., Irons, W.A., Samson, F.K., 1997. Monaural spectral contrast mechanism for neural sensitivity to sound direction in the medial geniculate body of the cat. J Neurophysiol 78(5), 2754–2771. 10.1152/jn.1997.78.5.2754.

Kim, D.O., Bishop, B., Kuwada, S., 2010. Acoustic cues for sound source distance and azimuth in rabbits, a racquetball and a rigid spherical model. J Assoc Res Otolaryngol 11(4), 541–557. 10.1007/s10162-010-0221-8.

Kim, D.O., Carney, L., Kuwada, S., 2020. Amplitude modulation transfer functions reveal opposing populations within both the inferior colliculus and medial geniculate body. J Neurophysiol 124(4), 1198–1215. 10.1152/jn.00279.2020.

Kistler, D.J., Wightman, F.L., 1992. A model of head-related transfer functions based on principal components analysis and minimum-phase reconstruction. J Acoust Soc Am 91(3), 1637–1647.

Koka, K., Jones, H.G., Thornton, J.L., Lupo, J.E., Tollin, D.J., 2011. Sound pressure transformations by the head and pinnae of the adult Chinchilla (Chinchilla lanigera). Hear Res 272(1-2), 135–147. 10.1016/j.heares.2010.10.007.

Koka, K., Read, H.L., Tollin, D.J., 2008. The acoustical cues to sound location in the rat: measurements of directional transfer functions. J Acoust Soc Am 123(6), 4297–4309. 10.1121/1.2916587.

Kuhn, G.F., 1977. Model for the interaural time differences in the azimuthal plane. J Acoust Soc Am 62(1), 157–167.

Kulkarni, A., Colburn, H.S., 1998. Role of spectral detail in sound-source localization. Nature 396(6713), 747–749. 10.1038/25526.

Kuwada, S., Kim, D.O., Koch, K.J., Abrams, K.S., Idrobo, F., Zahorik, P., Carney, L.H., 2015. Near-field discrimination of sound source distance in the rabbit. J Assoc Res Otolaryngol 16(2), 255–262. 10.1007/s10162-014-0505-5.

Langendijk, E.H., Bronkhorst, A.W., 2002. Contribution of spectral cues to human sound localization. J Acoust Soc Am 112(4), 1583–1596. 10.1121/1.1501901.

Lauer, A.M., Slee, S.J., May, B.J., 2011. Acoustic basis of directional acuity in laboratory mice. J Assoc Res Otolaryngol 12(5), 633–645. 10.1007/s10162-011-0279-y.

Maki, K., Furukawa, S., 2005. Acoustical cues for sound localization by the Mongolian gerbil, Meriones unguiculatus. J Acoust Soc Am 118(2), 872–886.

Mehrgardt, S., Mellert, V., 1977. Transformation characteristics of the external human ear. J Acoust Soc Am 61(6), 1567–1576. 10.1121/1.381470.

Middlebrooks, J.C., 1999. Individual differences in external-ear transfer functions reduced by scaling in frequency. J Acoust Soc Am 106(3 Pt 1), 1480–1492. 10.1121/1.427176.

Middlebrooks, J.C., Green, D.M., 1990. Directional dependence of interaural envelope delays. J Acoust Soc Am 87(5), 2149–2162.

Middlebrooks, J.C., Green, D.M., 1991. Sound localization by human listeners. Annu Rev Psychol 42, 135–159. 10.1146/annurev.ps.42.020191.001031.

Middlebrooks, J.C., Makous, J.C., Green, D.M., 1989. Directional sensitivity of sound-pressure levels in the human ear canal. J Acoust Soc Am 86(1), 89–108.

Musicant, A.D., Chan, J.C., Hind, J.E., 1990. Direction-dependent spectral properties of cat external ear: new data and cross-species comparisons. J Acoust Soc Am 87(2), 757–781. 10.1121/1.399545.

Neti, C., Young, E.D., Schneider, M.H., 1992. Neural network models of sound localization based on directional filtering by the pinna. J Acoust Soc Am 92(6), 3140–3156. 10.1121/1.404210.

Rabbitt, R.D., Holmes, M.H., 1988. Three-dimensional acoustic waves in the ear canal and their interaction with the tympanic membrane. J Acoust Soc Am 83(3), 1064–1080. 10.1121/1.396051.

Ramachandran, R., Davis, K.A., May, B.J., 1999. Single-unit responses in the inferior colliculus of decerebrate cats. I. Classification based on frequency response maps. J Neurophysiol 82(1), 152–163.

Rice, J.J., May, B.J., Spirou, G.A., Young, E.D., 1992. Pinna-based spectral cues for sound localization in cat. Hear Res 58(2), 132–152. 10.1016/0378-5955(92)90123-5.

Roth, G.L., Kochhar, R.K., Hind, J.E., 1980. Interaural time differences: implications regarding the neurophysiology of sound localization. J Acoust Soc Am 68(6), 1643–1651. 10.1121/1.385196.

Schnupp, J.W., Booth, J., King, A.J., 2003. Modeling individual differences in ferret external ear transfer functions. J Acoust Soc Am 113(4 Pt 1), 2021–2030. 10.1121/1.1547460.

Schroeder, M., 1970. Synthesis of low-peak-factor signals and binary sequences with low autocorrelation (Corresp.). IEEE Transactions on Information Theory 16(1), 85–89. 10.1109/TIT.1970.1054411.

Shaw, E.A., 1974. Transformation of sound pressure level from the free field to the eardrum in the horizontal plane. J Acoust Soc Am 56(6), 1848–1861. 10.1121/1.1903522.

Shaw, E.A., Teranishi, R., 1968. Sound pressure generated in an external-ear replica and real human ears by a nearby point source. J Acoust Soc Am 44(1), 240–249. 10.1121/1.1911059.

Slee, S.J., Young, E.D., 2010. Sound localization cues in the marmoset monkey. Hear Res 260(1-2), 96–108. 10.1016/j.heares.2009.12.001.

Wagner, J.D., Gelman, A., Hancock, K.E., Chung, Y., Delgutte, B., 2022. Rabbits use both spectral and temporal cues to discriminate the fundamental frequency of harmonic complexes with missing fundamentals. J Neurophysiol 127(1), 290–312. 10.1152/jn.00366.2021.

Wenzel, E.M., Arruda, M., Kistler, D.J., Wightman, F.L., 1993. Localization using nonindividualized head-related transfer functions. J Acoust Soc Am 94(1), 111–123. 10.1121/1.407089.

Wightman, F.L., Kistler, D.J., 1989. Headphone simulation of free-field listening. I: Stimulus synthesis. J Acoust Soc Am 85(2), 858–867. 10.1121/1.397557.

Xu, L., Middlebrooks, J.C., 2000. Individual differences in external-ear transfer functions of cats. J Acoust Soc Am 107(3), 1451–1459. 10.1121/1.428432.

Zakarauskas, P., Cynader, M.S., 1993. A computational theory of spectral cue localization. J Acoust Soc Am 94(3), 1323–1331.

Zhai, X., Khatami, F., Sadeghi, M., He, F., Read, H.L., Stevenson, I.H., Escabi, M.A., 2020. Distinct neural ensemble response statistics are associated with recognition and discrimination of natural sound textures. Proc Natl Acad Sci U S A 117(49), 31482–31493. 10.1073/pnas.2005644117.

Zhang, X., Heinz, M.G., Bruce, I.C., Carney, L.H., 2001. A phenomenological model for the responses of auditory-nerve fibers: I. Nonlinear tuning with compression and suppression. J Acoust Soc Am 109(2), 648–670. 10.1121/1.1336503.

